# Identification of microRNAs in the West Nile virus vector *Culex tarsalis*

**DOI:** 10.1101/2021.09.23.461553

**Authors:** Sultan Asad, Ahmed M. Mehdi, Sujit Pujari, Claudia Rüeckert, Gregory D. Ebel, Jason L. Rasgon

## Abstract

**Background:** microRNAs (miRNAs) represent a group of small non-coding RNAs that are crucial gene regulators of important biological functions including development and pathogen defense in most living organisms. Presently, there is a lack of availability of information regarding the miRNAs in the mosquito *Culex tarsalis*, which is one of the most important vectors of West Nile virus (WNV) in the United States. We used small RNA sequencing data and *in vitr*o and *in vivo* experiments to identify and validate the presence of a repertoire of miRNAs in *Cx. tarsalis* mosquitoes.

**Results:** Using bioinformatic approaches we analyzed small RNA sequencing data from the *Cx. tarsalis* CT embryonic cell line to discover 86 miRNAs. Consistent with other mosquitoes such as *Aedes albopictus* and *Cx. quinquefasciatus*, mi-184 was found to be the most abundant miRNA in *Cx tarsalis*. We also identified an additional 20 novel miRNAs from the recently sequenced *Cx. tarsalis* genome, for a total of 106 miRNAs identified in this study. The presence of selected miRNAs was biologically validated in both cell line and adult *Cx. tarsalis* mosquitoes using RT-qPCR and sequencing.

**Conclusions:** *Cx. tarsalis* is an important vector of many medically important pathogens including WNV and Western Equine encephalitis. Here we report a detailed insight into the miRNA population in *Cx. tarsalis* mosquitoes. These results will open new avenues of research deciphering the role of miRNAs in different *Cx. tarsalis* biological events such as development, metabolism, immunity and pathogen infection.

## Background

Recent reports of the involvement of miRNAs in insect immunity and host-pathogen interactions have highlighted the need for investigating miRNAs in mosquitoes that have significant impacts on global health. miRNAs are small non-coding RNAs ranging from 19-25 nucleotides in length which regulate post transcriptional gene expression [1, 2]. In the animal kingdom, miRNAs regulate gene expression by binding imperfectly to the mRNAs either at the 3’-untranslated regions (3’-UTR), 5’-untranslated regions (5’-UTR), or coding regions. Though the 2-8 nucleotides at the 5’ end of the miRNA (“seed region”) perfect complementarity to the target transcript is integral for miRNA function, the rest of the miRNA sequence can have mismatches and bulges [1, 3, 4]. miRNAs are transcribed inside the nucleus into a primary miRNA transcript which is processed into a pre-miRNA by Drosha and Pasha. The pre-miRNA is then exported to the cytoplasm with the help of exportin proteins. Once in cytoplasm, the pre-miRNA is processed by Dicer I into the ∼22 nucleotide mature miRNA. Once maturation is completed the mature miRNA is loaded into the microRNA induced silencing complex (miRISC) where it becomes single-stranded. The single-stranded miRNA-miRISC complex then targets the mRNA with sequence similarity leading to target mRNA degradation or modulation of protein expression [5, 6]. miRNAs are critical for a variety of cellular processes including development [7], immunity [8] and pathogen response [9, 10], which make them an interesting choice to investigate in the host pathogen interactions of vector mosquitoes and the medically important pathogens they carry.

More than 3000 mosquito species are present worldwide, but many important mosquito species are underrepresented in miRbase (a database of all miRNAs reported from different species) [11]. Recent advances in sequencing platforms and bioinformatic approaches have allowed us to successfully identify the miRNAs from species whose genomes are not yet properly sequenced, assembled and annotated.

*Cx. tarsalis* is a neglected yet important vector of many medically important viruses such as WNV [12-14], West Equine Encephalitis (WEEV) [15], St. Louis Encephalitis (SLEV) [16] and Cache Valley virus (CVV) [17] in North America. Despite many studies focusing on the ecology, feeding behavior and vector competence of this mosquito, there is little information regarding miRNAs in *Cx. tarsalis*. Here we have mined and analyzed publicly available small RNA sequencing data of the *Cx. tarsalis CT* cell line to identify 86 high confidence miRNAs that are present in *Cx tarsalis* mosquitoes. Ten randomly chosen miRNAs were confirmed both *in-vitro* and *in-vivo* in mosquitoes.

## Results and Discussion

### Data Used in this study

We accessed publicly available small RNA data generated from the *Cx. tarsalis* CT cell line and passed it through our in-house pipeline (Figure 1) to identify miRNAs, using CLC genomic work bench 20. SRA read files described in Figure 2A were directly downloaded into CLC genomics workbench. All reads were trimmed by removing the RA3 adapter (TGGAATTCTCGGGTGCCAAGGG) sequences from the 3’ end of raw sequencing reads. Any reads with low quality scores or that did not have an adapter sequence were excluded. After removal of the RA3 adapter sequences we saw a cluster of RNA sequences ranging from 21-24 nucleotides with the highest peak at 22 nucleotides which is the most abundant size of mosquito miRNAs (Figure 2B). In order to further examine the profile of the trimmed RNA reads, we used the web-based server sRNA toolbox [18] to determine that approximately 70 percent of the reads were mappable to miRbase (Figure 2C).

**Figure 1.**
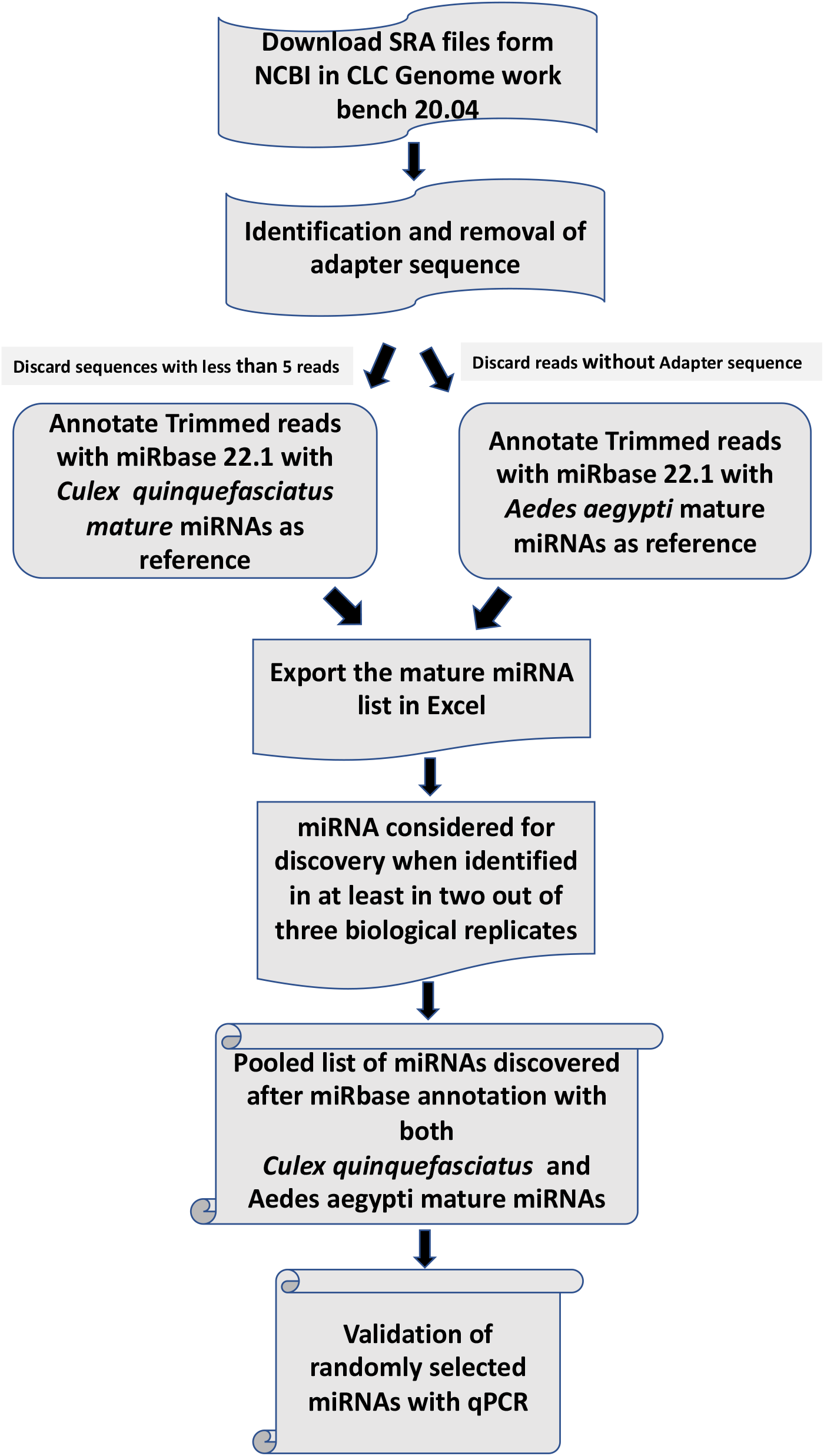
Schematic diagram of miRNA discovery pipeline. Diagram showing use of an integrated approach of bioinformatics and wet laboratory experiments to identify *Cx. tarsalis* miRNAs.

**Figure 2.**
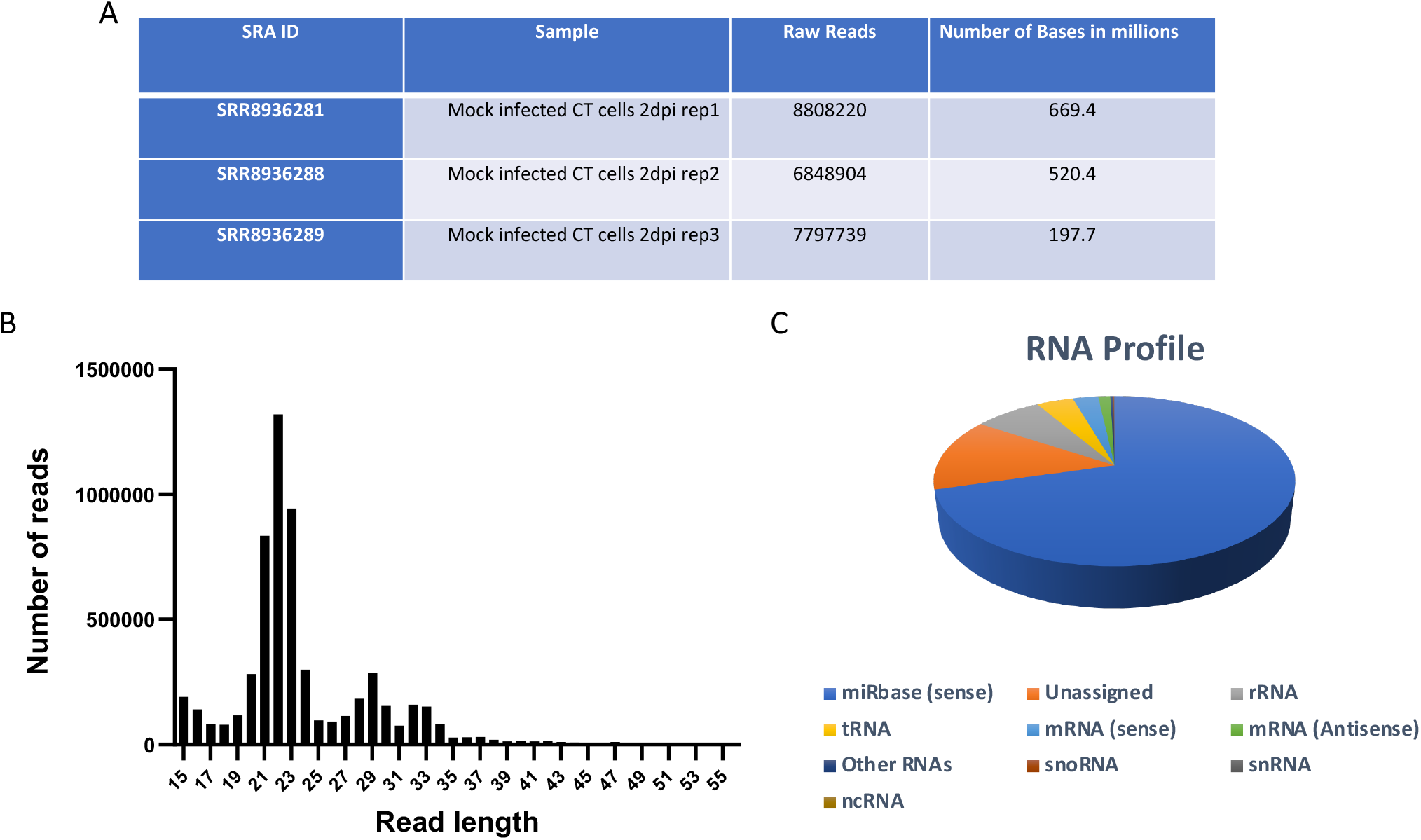
Small RNA sequencing data. A*) Culex tarsalis* small RNA data used in this study. B) Read length distribution after the adapter removal showed enrichment of reads around 21-24 nucleotides confirming the successful removal of adapter sequences. C) RNA profile of reads after adapter removal showing around 70 percent of read mapped to miRbase.

### Discovery of miRNAs in *Cx tarsalis*

miRNAs have been demonstrated to play a pivotal role in insect development [7, 19], reproduction [20, 21], metabolism and longevity [22, 23], insecticide resistance [24, 25], immunity, and host-pathogen interactions [26, 27], and have been shown to govern processes relevant to public health in various mosquito species [28-30]. Using CLC genomic workbench 20 we annotated and mapped trimmed small RNA reads from 3 biological replicates from the *Cx. tarsalis* CT cell line to mature miRNAs of both *Ae. aegypti* and *Cx. quinquefasciatus* using default parameters. miRNAs discovered were considered to be true hits only if they were present in at least two out of three biological replicates. Our analysis found 60 and 84 mature miRNAs in *Cx. tarsalis* using *Cx. quinquefasciatus* (Figure 3A) and *Ae. aegypti* (Figure 3B) mature miRNAs as reference, respectively, with a 58 mature miRNA overlap using Jvenn online server (figure 3C, Supplementary table 1) [31]. Finally, we successfully identified a total of 86 high confidence miRNAs in *Cx. tarsalis* mosquitoes (Table 1).

**Figure 3.**
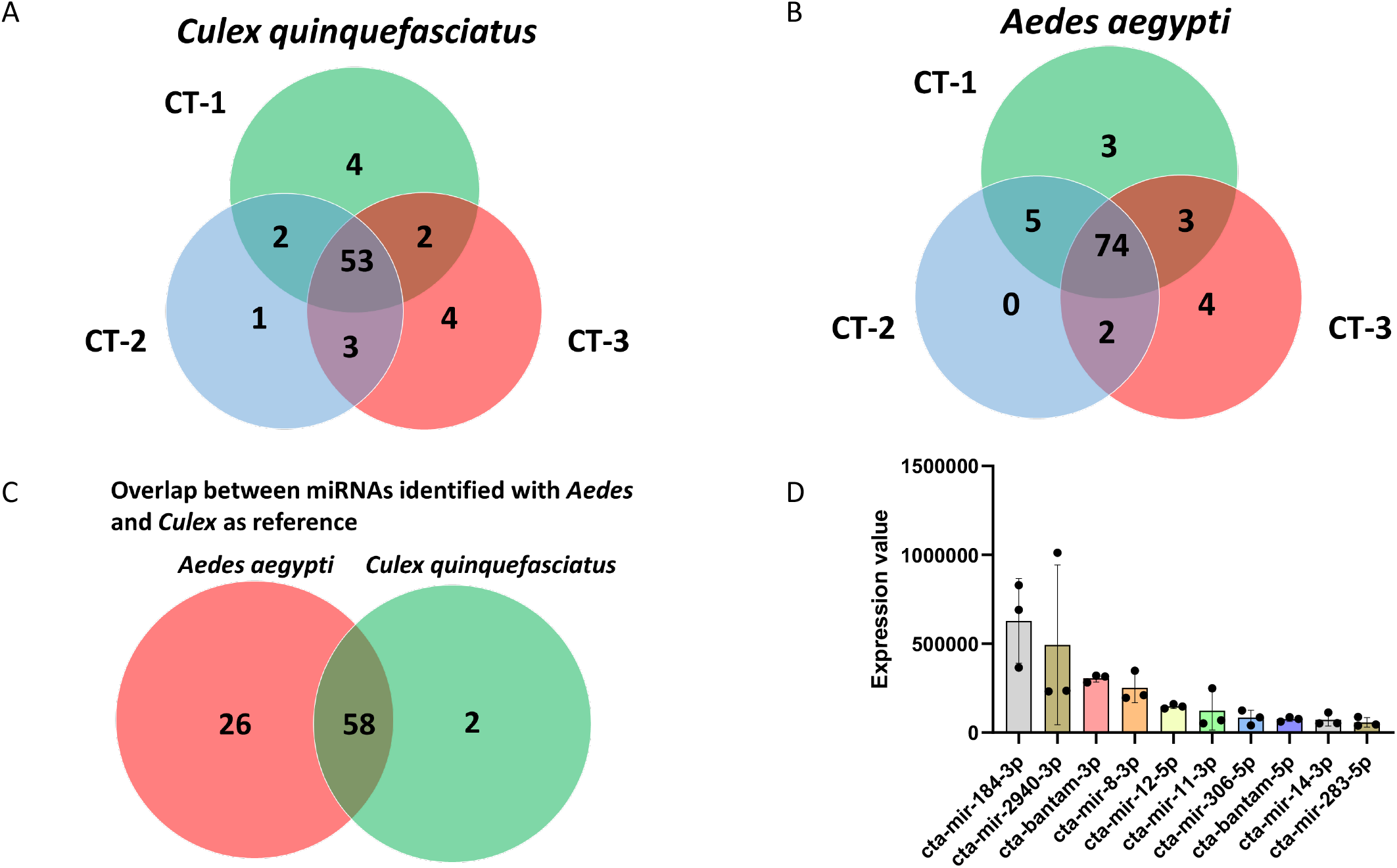
Identification of *Cx tarsalis* miRNAs. A) Venn diagram of miRNAs identified across 3 biological replicates. B) Venn diagram showing common miRNAs identified within 3 biological replicates. C) Overlap of miRNAs identified with *Ae. aegypti* and *Cx. quinquefasciatus* annotation. D) 10 most abundant miRNAs in *Cx. tarsalis* RNA samples, each bar represents average normalized expression value of individual miRNA, error bars represent the standard deviation, and the dots shows the values of individual biological replicates.

**Table 1.**
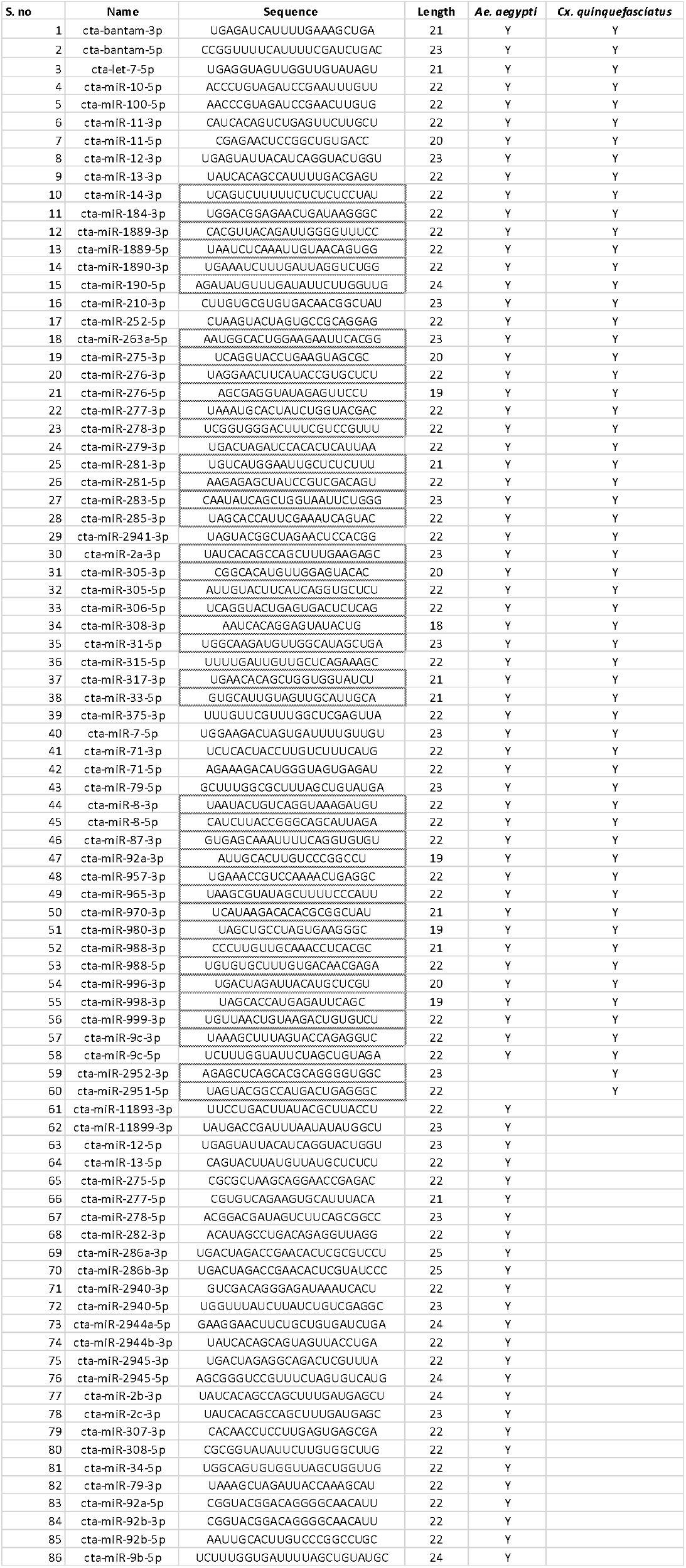
86 miRNAs identified by homology to *Ae. aegypti* and *Cx. quinquefasciatus*.

### Majority of *Cx. tarsalis* miRNAs arise from the 3’ arm and sequence variations occur mostly at 3’ arm of mature miRNAs

Mature miRNAs are formed as a result of a Dicer-mediated processing of precursor miRNA (pre-miRNA) having two arms/strands named 5p and 3p, leading to selective loading and retention of either 3p or 5p in the miRISC complex to become functionally active, while the other strand is cleaved out of the complex and degraded [5]. This strand selection has been demonstrated to be influenced by the thermodynamic instability of the duplex, 5’ end starting nucleotides, and miRNA duplex length [32-34]. In our results we were able to detect mature miRNA that arose from 5p, 3p and both 5p and 3p of pre-miRNAs. Our data showed that 49 out of 86 miRNAs identified in this study were derived from the 3’ arm of the pre-miRNA which indicates that in the *Cx. tarsalis* CT cell line, there may be a preference toward 3p arm processing of miRNAs (Figure 4A). However, this finding needs further experimental validation.

**Figure 4.**
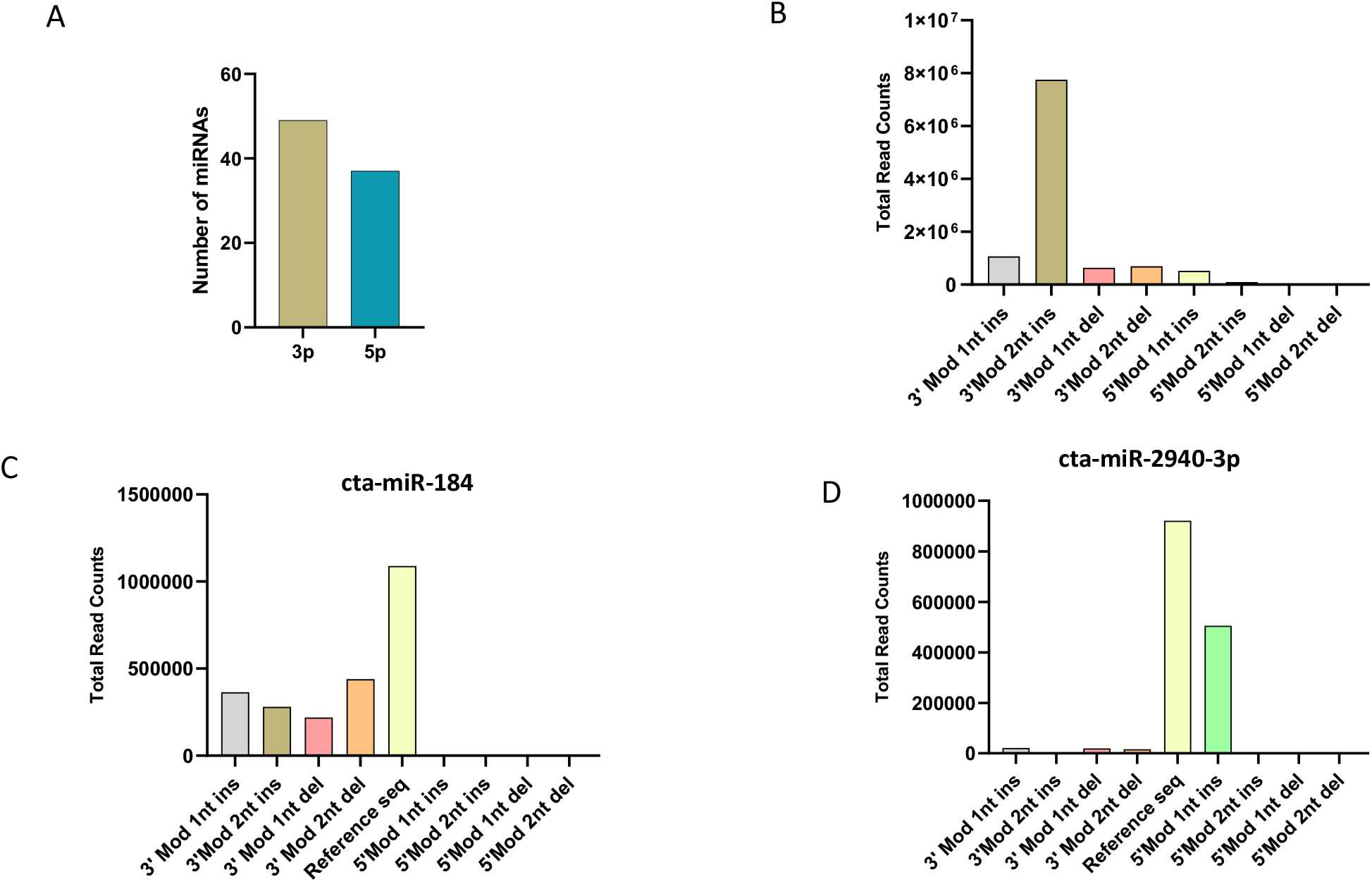
*Cx. tarsalis* miRNA mostly arise from 3p arm having most of sequence variation at 3’end of miRNA. A) Number of miRNAs processed from either 3p or 5p arm of pre-miRNA. B) Overall sequence variations at 3’ or 5’ of mature miRNA sequences. C) Sequence variation found in cta-miR-184 showing most of the variations at the 3’ end of the mature miRNA. D) Sequence variation found in cta-miR-2940-3p showing most of the variations at the 5’ end of the mature miRNA.

In each of the data libraries used in this study, adapter trimmed small RNA reads mapped to the reference sequences of mature miRNAs from miRbase showed variability. These variants with slight difference in length and sequence are commonly known as isomiRs. These isomiRs are usually produced due to imperfect processing/cleavage by Drosha or Dicer [35-37]. IsomiRs were long considered as an artifact, but recent advances in deep sequencing and computational algorithms have identified isomiRs in many species [38-40], including in *Ae. aegypti* [41]. IsomiRs are classified into three main categories: 5’ isomiRs, 3’ isomiRs, and polymorphic isomiRs with 5’ and 3’ isomiRs subclassified according to the nucleotide’s insertions and deletions [42]. To find out and quantify the miRNA isomiRs, we used CLC Genomic workbench 20 to quantify the isomiRs based on nucleotide additions, insertions and substitutions. We focused on insertions and deletions of nucleotides at 3’ or 5’ of mature miRNAs and limited our discovery to 2 nucleotide addition or deletion at the either end. IsomiRs that were present in all three biological replicates were used for analysis (Supplementary table 2). Our results show most modifications occurring at the 3’ end of the mature miRNAs, in particularly 2 nucleotide insertion followed by the 1 nucleotide insertion (Figure 4B). Our results agree with previous studies that highlighted majority of sequence variations leading to isomiRs formation are at the 3’ end of mature miRNAs [43]. Although the 5’ variations are rare, they have more evolutionary and functional importance, as most of the 5’ variations lead to a shift in the seed region that might affect regulatory ability of the miRNA [44]. Expanded analysis revealed that cta-miR-184 showed a total of 240 different types of isomiRs, followed by cta-miR-2940-3p that showed 161 different types of isomiRs. Although both cta-miR-184 and cta-miR-2940-3p showed the highest amount of isomiRs formation, there was a striking difference in the modification pattern as the former showed the majority of sequence variations at the 3’ end (Figure 4C) while the later showed the highest amount of sequence variations at the 5’ end (Figure 4D). These results warrant further investigation into the role of isomiRs in mosquito physiology and host pathogen interactions, for example in the case of WNV infections.

**Table 2.**
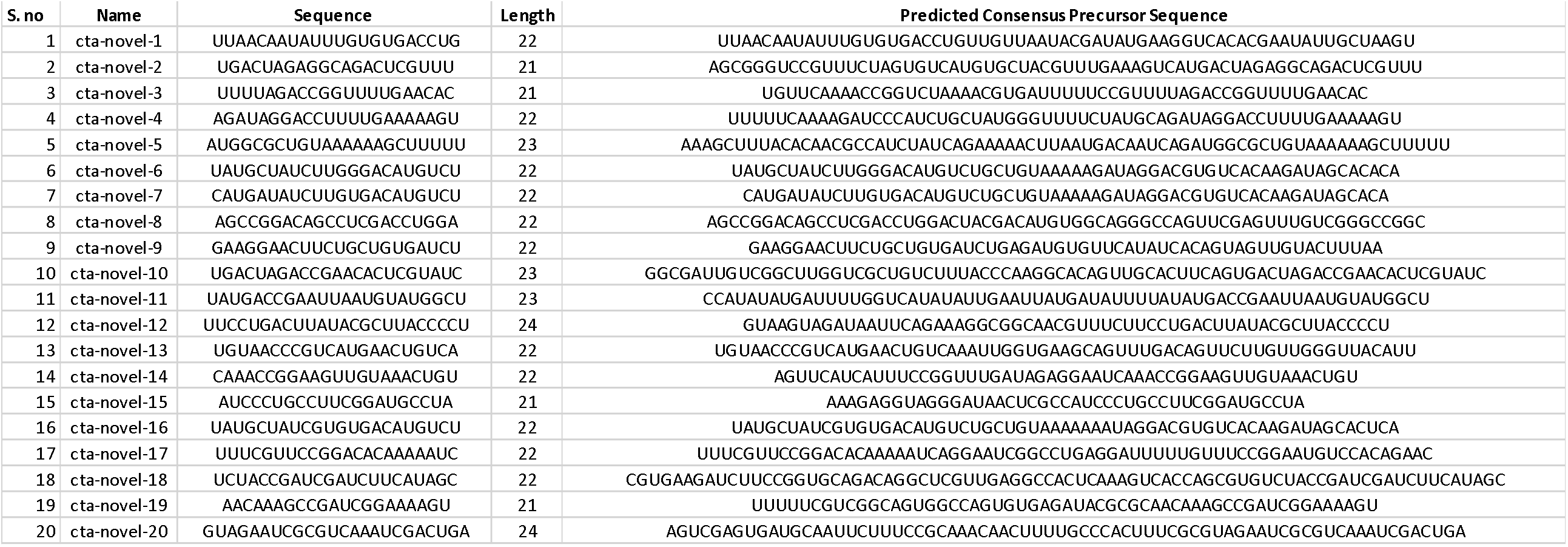
Novel miRNAs identified from the *Cx. tarsalis* genome.

### Validation of *Cx. tarsalis* miRNAs

We randomly selected 10 miRNAs: cta-miR-317-3p, cta-miR-7, cta-miR-999, cta-miR-71-3p, cta-miR-33, cta-miR-2940-5p, cta-miR-998, cta-miR-92b-3p, cta-miR-2951-5p and cta-miR-2945 for validation both *in vitro* (CT cell line) and *in-vivo* in 7 day old sugar fed female *Cx. tarsalis* mosquitoes (Figure 5A, B). All qPCR amplicons were cloned into pJET and mature miRNA sequences confirmed through Sanger sequencing (Supplementary Figure 1).

**Figure 5.**
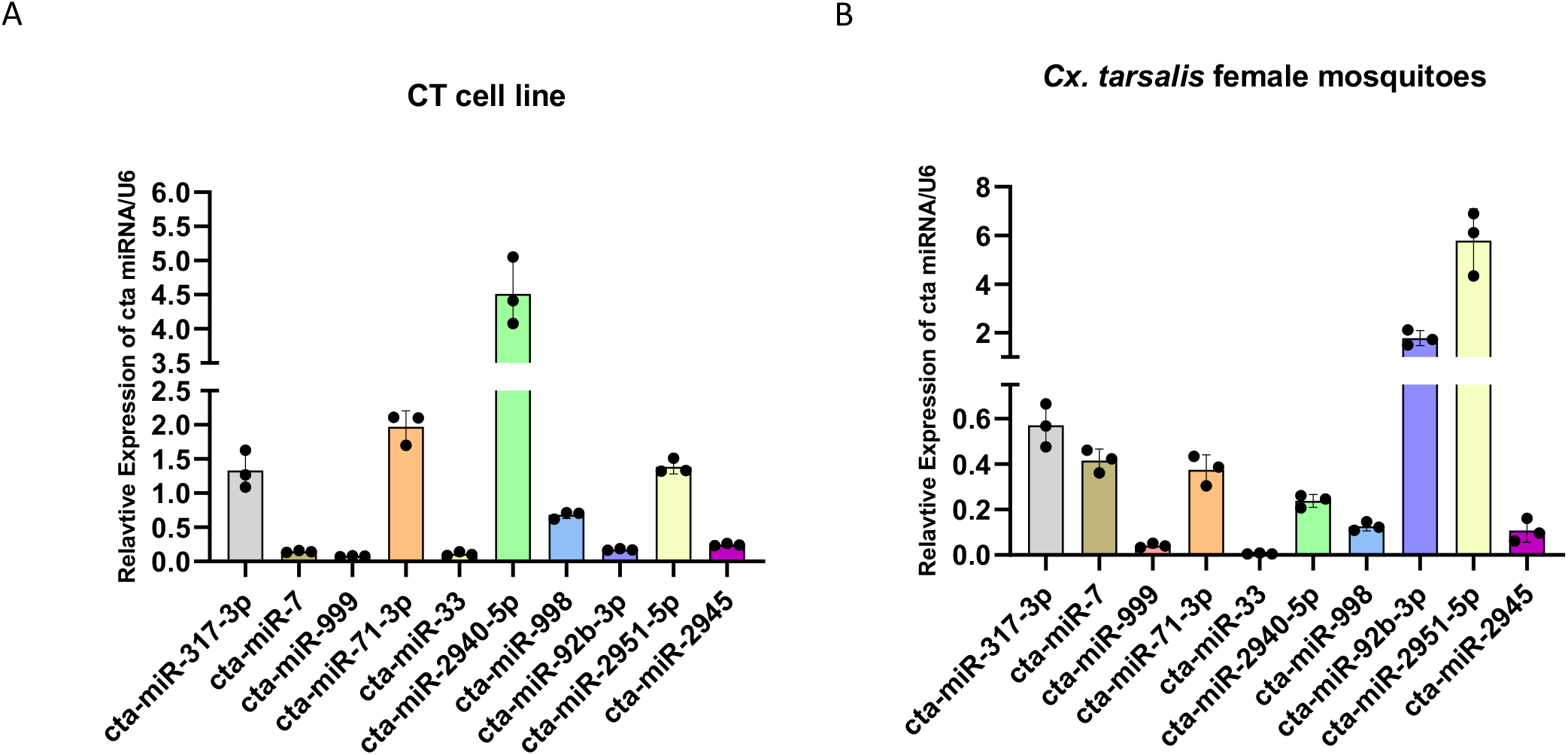
Validation of *Cx. tarsalis* miRNAs. Primer extension-based confirmation of 10 randomly selected miRNAs with RT-qPCR using 3 biological replicates. U6 was used as reference for relative quantification. Graphs showing relative expression of cta-miRNAs/U6 in A) CT cell line B) *Cx. tarsalis* female mosquitoes. Error bars represent standard deviation and dots represent individual biological replicate values. Each biological replicate is made up of 5 *Cx. tarsalis* sugar fed 7 days old females.

### Prediction and validation of novel miRNAs from the *Cx. tarsalis* genome

During the preparation of this manuscript, the *Cx. tarsalis* genome was published [45], which we used to identify novel miRNAs (present in all 3 replicates) that could be mapped to the *Cx. tarsalis* genome. Our results identified 20 novel miRNAs in *Cx. tarsalis* mosquitoes (Table 2). RNAfold (rna.tbi.univie.ac.at/cgi-bin/RNAWebSuite/RNAfold.cgi) was used to determine the secondary structure of the predicted pre-miRNA sequences (Figure 6). Primers were designed for 12 randomly selected novel miRNAs and validated through RT-qPCR in both the CT cell line and *Cx. tarsalis* mosquitoes. Our results confirm that all 12 novel selected miRNAs were expressed in CT cells (Figure 7A), but in mosquitoes we were only able to detect 8 out of 12 (Figure 7B), likely reflecting physiological differences between cultured cells and live mosquitoes.

**Figure 6.**
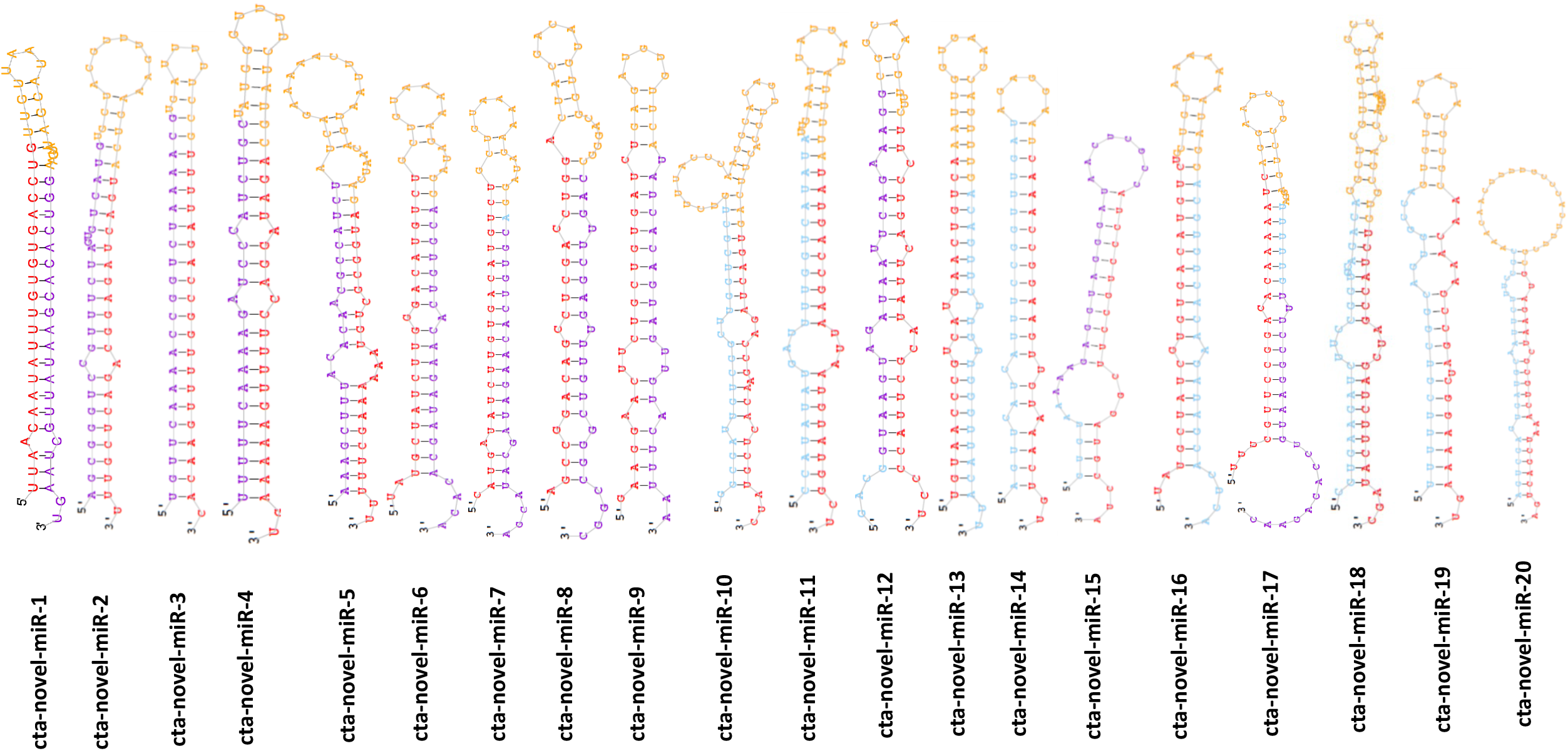
Prediction of secondary structure of novel miRNAs. Stem-loop structures of predicted novel pre-miRNAs. Mature novel miRNAs are highlighted in red while their corresponding strand are highlighted in blue/magenta.

**Figure 7.**
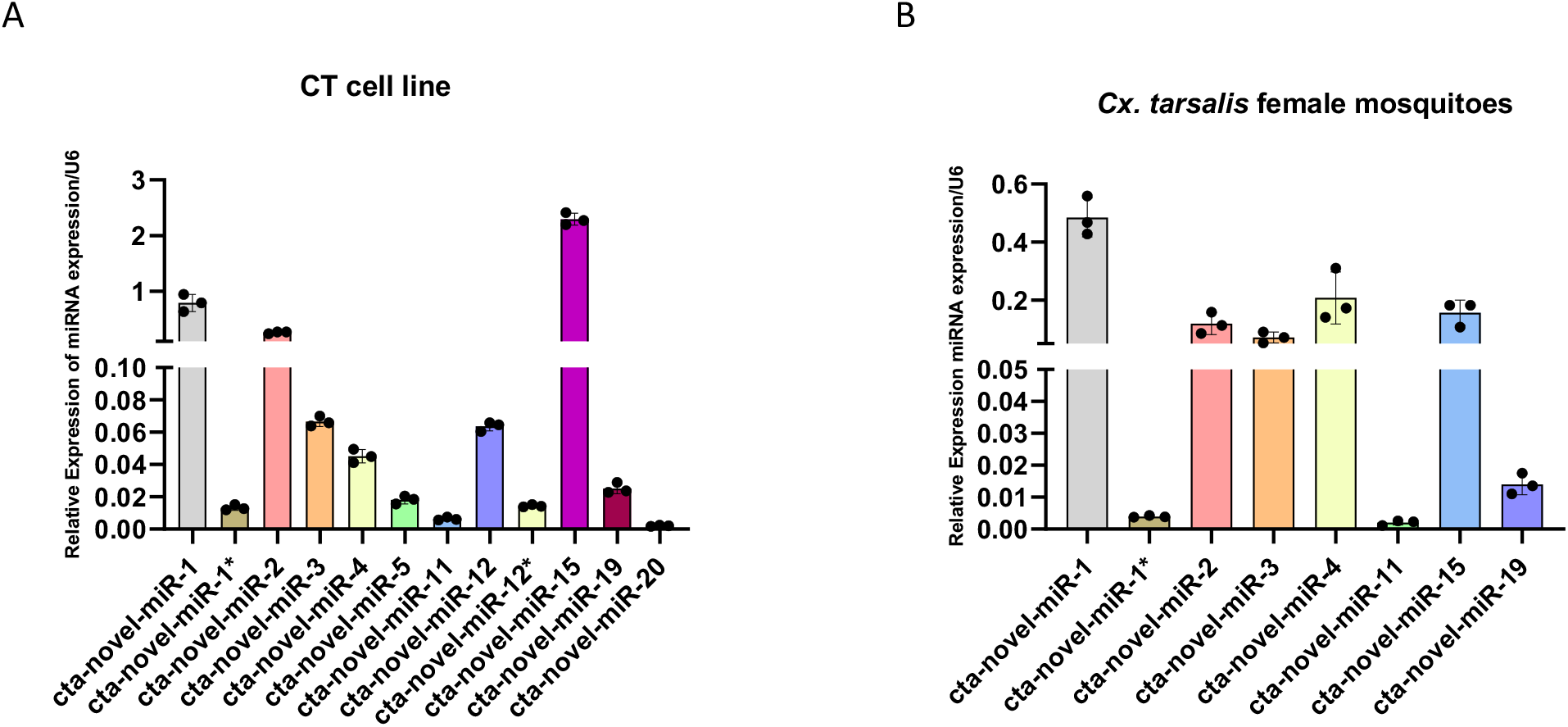
Validation of novel *Cx tarsalis* miRNAs. RT-qPCR based amplification of novel miRNAs in A) CT cells and B) *Cx. tarsalis* female mosquitoes. Error bars represent standard deviation and dots represent individual biological replicate values. Each biological replicate is made up of 5 sugar fed *Cx. tarsalis* 7 day old females.

## Conclusions

Our study has identified a total of 106 miRNAs, including 86 already reported in other mosquitoes, and 20 novel miRNAs that are present in the *Cx. tarsalis* genome. These results will lay the foundation for functional studies and open new avenues for research in *Cx. tarsalis* biology and pathogen transmission.

## Methods

### Small RNA libraries and data analysis

To identify *Cx. tarsalis* miRNAs we have used publicly available small RNA data of *Cx. tarsalis* CT cell line [46] from National Center for Biotechnology Information Sequence Read archive. A total of 3 libraries were downloaded. All libraries were generated using the TruSeq Small RNA Sample Prep Kit (Illumina) and sequenced on the Illumina HiSeq 2000 platform. The data were analyzed using CLC genome workbench 20. Libraries were trimmed of adapter sequences and the trimmed libraries were mapped to known miRNAs and miRNAs star strands of *Ae. aegypti* and *Cx. quinquefasciatus*. For novel miRNA prediction the miRDeep2 pipeline was used [47].

### Mosquitoes and cell lines

*Cx. tarsalis* mosquitoes (YOLO strain) were reared at 26 °C with 40-50% relative humidity and a 16:8 hr light and dark cycle. The mosquitoes were provided with 10% sucrose solution *ad libitum*. The *Cx. tarsalis* cell line CT (generously provided by Dr Aaron Brault, CDC) was maintained at 28 °C with 0.5% CO_2_ in Schneider medium supplemented with 10% of fetal bovine serum (GIBCO) and 1% antibiotics (GIBCO). Cells were passaged once per week.

### RNA extractions and miRNA RT-qPCR

Qiazol was used to extract total RNA from both cells and mosquitoes following the manufacturer’s suggested protocol. For mosquito RNA samples, 5 *Cx. tarsalis* sugar fed females (7 days old) were pooled per biological replicate. RNA was quantified using a Nanodrop and 2ug of total RNA was used to synthesize miRNA cDNA using the miScript RT kit following manufacturer’s suggested protocol. cDNA was 10 times diluted and approximately 18ng of cDNA was used to validate miRNAs identified in this study using the miScript qPCR kit according to manufacturer’s suggested protocol. All qPCR reactions were performed with 3 biological replicates and 2-3 technical replicates. U6 RNA was used as internal control for RT-qPCR. CT values were imported into Qiagen qPCR data analysis excel template to get normalized expression values.

### Cloning and sequencing of miRNA amplicons

qPCR amplicons were seperated using 1% agarose gel electrophoresis. Single amplicon bands were cut using a sterile surgical blade and DNA was extracted with the Zymo Gel extraction kit according to manufacturer’s suggested protocol. Eluted amplicons were directly cloned in to pJET1.2/Blunt vector according to manufacturer’s suggested protocol. Positive clones were sent to GENEWIZ for Sanger sequencing. Sequencing data was analyzed with CLC genomic workbench 20.

## Acknowledgements

This work was supported by NIH grants R01AI15025, R01AI128201, and R01AI116636 to JLR. We are thankful to Dr Aaron C. Brault from CDC for generously providing the CT cell line.

