## Supplementary Figure 1 for "Identification of microRNAs in the West Nile virus vector *Culex tarsalis*"

- cta-mir-317-3p: TGAACACAGCTGGTGGTATCT

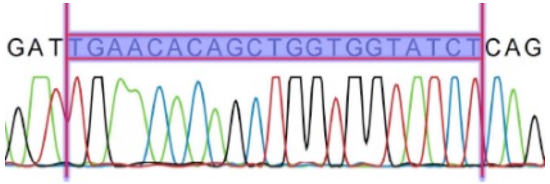


- cta-mir-7: TGGAAGACTAGTGATTTTGTTGT

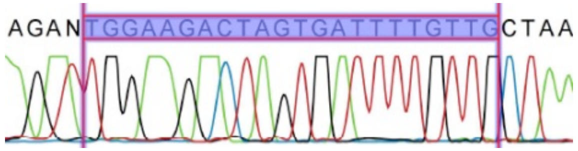


- cta-mir-999: TGTTAACTGTAAGACTGTGTCT

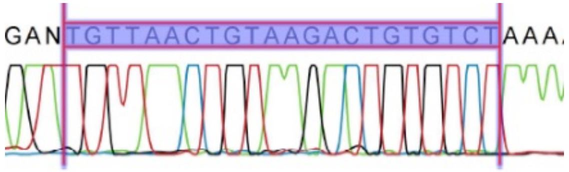


- cta-mir-71-3p: TCTCACTACCTTGTCTTTCATG

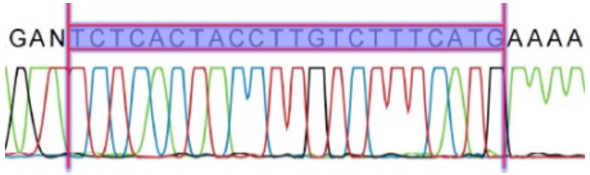


- cta-mir-33: GTGCATTGTAGTTGCATTGCA

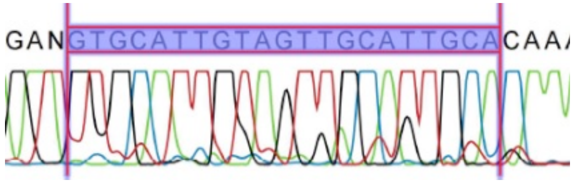

**Supplementary figure 1**

- cta-mir-2940-5p: TGGTTTATCTTATCTGTCGAGGC

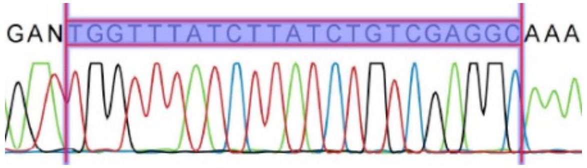


- cta-mir-998: TAGCACCATGAGATTCAGC

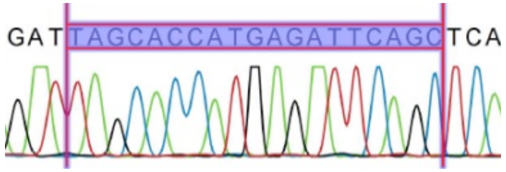


- cta-mir-92b-3p: AATTGCACTTGTCCCGGCCTGC

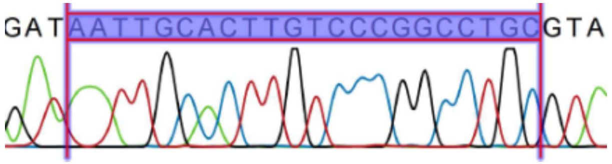


- cta-mir-2951-5p: AGAGCTCAGCACGCAGGGGTGGC

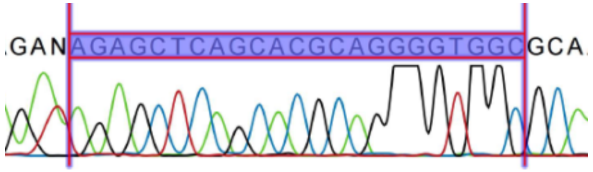


- cta-mir-2945-3p: TGACTAGAGGCAGACTCGTTTA

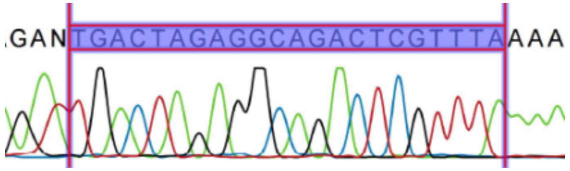
